# Structural basis for inhibition of human primase by arabinofuranosyl nucleoside analogues Fludarabine and Vidarabine

**DOI:** 10.1101/605279

**Authors:** Sandro Holzer, Neil J. Rzechorzek, Isobel R. Short, Michael Jenkyn-Bedford, Luca Pellegrini, Mairi L. Kilkenny

**Affiliations:** Department of Biochemistry, University of Cambridge, Cambridge, CB2 1GA, UK

## Abstract

Nucleoside analogues are widely used in clinical practice as chemotherapy drugs. Arabinose nucleoside derivatives such as Fludarabine are effective in the treatment of patients with acute and chronic leukemias and non-Hodgkin lymphomas. Although nucleoside analogues are generally known to function by inhibiting DNA synthesis in rapidly proliferating cells, the identity of their *in vivo* targets and mechanism of action are often not known in molecular detail. Here we provide a structural basis for inhibition by arabinose nucleotides of human primase, the DNA-dependent RNA polymerase responsible for initiation of DNA synthesis in DNA replication. Our data suggest ways in which the chemical structure of Fludarabine could be modified to improve its specificity and affinity towards primase, possibly leading to less toxic and more effective therapeutic agents.

## INTRODUCTION

Prior to cell division, cells must accurately duplicate their genetic material to ensure that both daughter cells contain a full complement of genes. The process of DNA replication is carried out by a large and dynamic macromolecular assembly known as the replisome, which co-ordinates unwinding of the DNA duplex with DNA synthesis of both leading and lagging strands (Pellegrini and Costa, 2016). Because replicative DNA polymerases cannot initiate the synthesis of new DNA, they rely on a DNA-dependent RNA polymerase known as primase to produce short RNA oligonucleotides that act as primers (Frick and Richardson, 2002). Primase therefore plays an essential role in DNA replication, priming the synthesis of both the leading and lagging strand.

Human primase is a heterodimeric enzyme comprising two subunits: Pri1 (49.9 kDa, also known as PriS or p49) and Pri2 (58.8 kDa, also known as PriL or p58), encoded by the *PRIM1* and *PRIM2* genes, respectively (Lucchini et al., 1987). *PRIM1* maps to a region of chromosome 12 that is amplified in numerous tumour types (Cloutier et al., 1997; Yotov et al., 1999). In addition, elevated *PRIM1* expression has been observed in breast tumour tissues, correlating with poorer patient outcomes (Lee et al., 2018). It has therefore been suggested that primase may represent an effective target for anti-cancer therapy. More recently, a CRISPR genome-wide dropout screen identified *PRIM1* as an essential gene in all 7 cancer cell lines tested (Tzelepis et al., 2016). Synthetic lethality has been demonstrated between *PRIM1* and *ATR* across a range of cancer cell lines, leading to renewed interest in primase inhibitors as anti-cancer therapeutics in ATR-deficient tumours (Hocke et al., 2016; Job et al., 2018).

Many anti-cancer agents in clinical use interfere with tumour growth by inhibiting DNA replication. A subset of these replication inhibitors, known as nucleoside analogues, comprise a series of pyrimidine and purine nucleoside antimetabolites that are widely used in the treatment of haematological malignancies and solid tumours (Plunkett and Saunders, 1991; Jordheim et al., 2013). These compounds include Fludarabine (2F-araAMP), Vidarabine (araA), Cytarabine (araC), Cladribine (2Cl-dA), Gemcitabine (2′,2′-diF-dC) and Clofarabine (2Cl-2′F-aradA). Upon cellular uptake, these analogues are biologically activated by 5′-triphosphorylation. They subsequently elicit their effects by directly inhibiting intracellular enzymes and/or by retarding or terminating nucleic acid synthesis as they are incorporated into nascent DNA and RNA strands (Jordheim et al., 2013; Sandoval et al., 2002).

Vidarabine triphosphate (Vidarabine-TP) and Fludarabine triphosphate (Fludarabine-TP) are both ATP analogues in which the 2′-hydroxyl is in the *arabino (ara)* rather than the *ribo* configuration (Fig. 1A). Vidarabine, while no longer used as a cancer treatment due to its rapid deamination *in vivo*, is nonetheless effective as an antiviral agent against *herpes simplex virus* and *varicella zoster virus* infections (De Clercq and Li, 2016). Fludarabine, which is more resistant to deamination, is widely used as a chemotherapeutic agent to treat B-cell chronic lymphocytic leukemia (CLL), acute myeloid leukaemia (AML) and some types of non-Hodgkin lymphoma (Chun et al., 1991; Tamamyan et al., 2017). However, treatment is often associated with thrombocytopenia, anemia, neutropenia and profound lymphopenia, thereby increasing the risk of opportunistic infections (Lukenbill and Kalaycio, 2013; Fidias et al., 1996). In fact, one of the major problems with the therapeutic use of current nucleoside analogues is dose-limiting toxicity due to their non-selective nature, with potential targets including DNA and RNA polymerases, ribonucleotide reductase and DNA ligase (Parker et al., 1988; Tseng et al., 1982; White et al., 1982; Yang et al., 1992; Huang et al., 1990; Wisitpitthaya et al., 2016; Huang and Plunkett, 1991). Vidarabine-TP and Fludarabine-TP are potent inhibitors of DNA replication *in vivo* (Plunkett et al., 1980), and have been shown to inhibit both replicative DNA polymerases and primase *in vitro* (Dicioccio and Srivastava, 1977; Yoshida et al., 1985; Parker and Cheng, 1986). Primase incorporates *ara* nucleotides into primers more efficiently than normal ribonucleotides (Catapano et al., 1993; Harrington and Perrino, 1995; Kuchta et al., 1992), and it has been suggested that primase inhibition may be one of the primary mechanisms for Fludarabine cytotoxicity in tumour cells (Kuchta and Willhelm, 1991; Catapano et al., 1993; Catapano et al., 1991). However, deciphering which of the many nucleotide-binding enzymes are the primary targets of the *ara* nucleotides *in vivo* has proved difficult, with the result that their mechanism of cytotoxicity is still unclear. A drug that selectively inhibits primase without significant off-target effects could be of significant therapeutic value, and may be less toxic to non-dividing cells (Moore et al., 2002). Importantly, recent structural information has made the design of high affinity primase inhibitors a more realistic prospect (Kilkenny et al., 2013; Vaithiyalingam et al., 2014). Here we test the effect of a range of chemotherapeutic nucleotide analogues on primase activity. We also present the x-ray crystal structures of human Pri1 bound to Vidarabine-TP and Fludarabine-TP, thereby elucidating the mode of binding of arabinofuranosyl nucleotides to the catalytic subunit of primase, and explaining the reported preference of primase for these nucleotides. We propose that these *ara* nucleotides represent an interesting starting point for the structure-based drug design of specific primase inhibitors.

**Figure 1:**
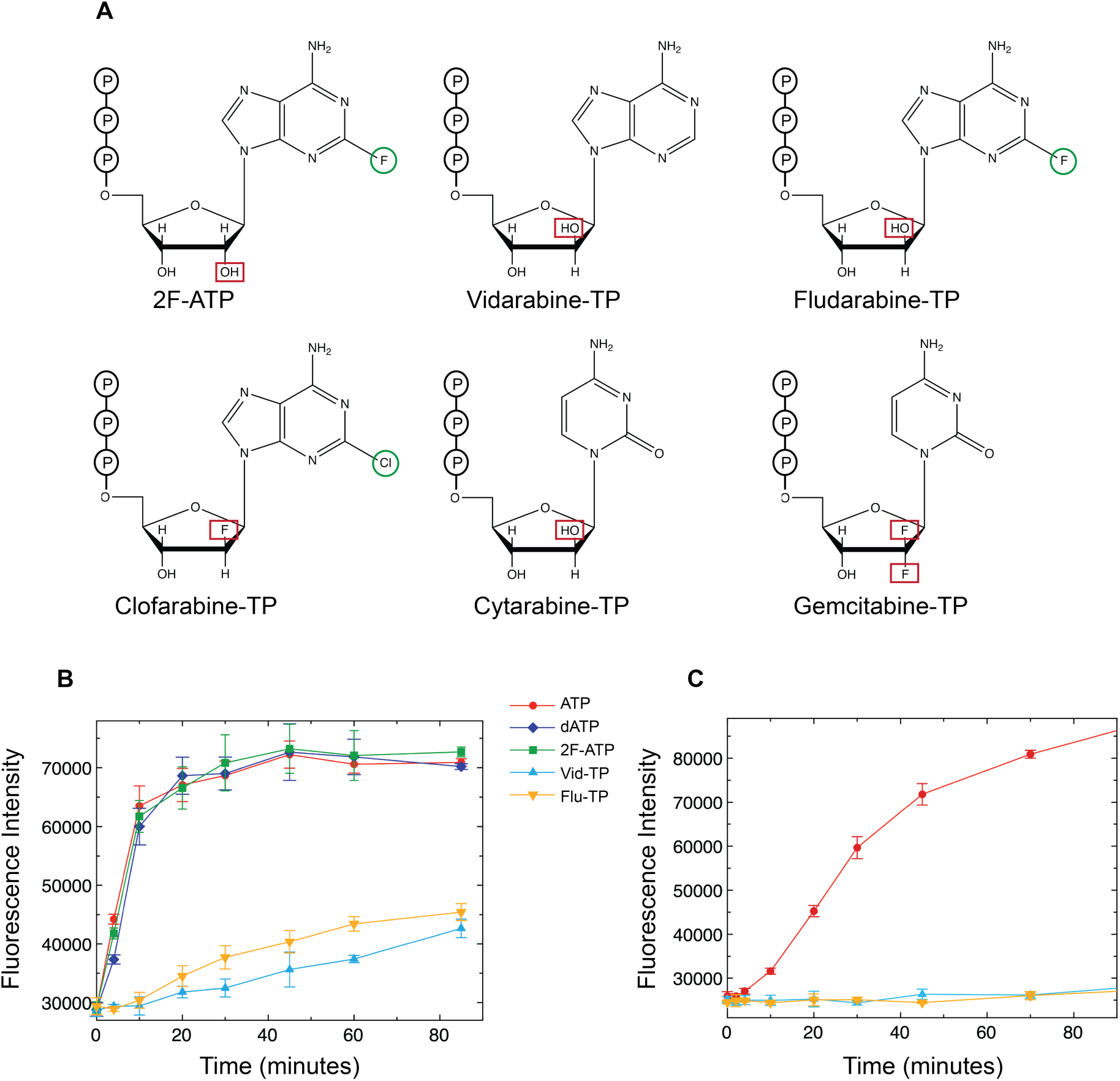
The *ara* nucleotides inhibit RNA primer synthesis by human primase. (a) Chemical structures of the nucleotide analogues used in this study. Structures were generated using ChemDraw 18.0. (b) Fluorescence-based RNA primer synthesis assay on a ssDNA template (5′-GTTGTCCATTATGTCCTA CCTCGTGCTCCT) in the presence of Mn^2+^ ions and equimolar concentrations of ribonucleotides (20 µM each rNTP) and the indicated nucleotide analogue (20 µM). (c) As in (b) but with Mn^2+^ ions replaced by Mg^2+^ ions. Each data point represents the mean ± s.d. (n=3). Curves are coloured as follows: ATP (red), dATP (navy), 2F-ATP (green), Vidarabine-TP (light blue), Fludarabine-TP (orange).

## RESULTS

### Inhibition of RNA primer synthesis by Vidarabine-TP and Fludarabine-TP

To confirm that *ara* nucleotides are effective inhibitors of human primase, we analysed their effect on RNA primer synthesis. A fluorescence-based RNA primer synthesis assay (Koepsell et al., 2005) revealed strong inhibition of primase activity by both Vidarabine-TP and Fludarabine-TP (Fig. 1B, C). While similar levels of inhibition were observed for both compounds, no inhibition was observed in the presence of 2F-ATP, confirming that it is the arabinofuranosyl moiety that is important for mediating the inhibitory effect of these nucleotide analogues. Inhibition occurred irrespective of whether the divalent metal was Mn^2+^ (Fig. 1B) or Mg^2+^ (Fig. 1C). While both divalent metals support robust primer synthesis *in vitro*, Mn^2+^ has been reported to enhance the binding of nucleotides to human primase, but also to reduce the fidelity of various polymerases including primase (Kirk and Kuchta, 1999; Copeland et al., 1993; Vaithiyalingam et al., 2014).

These results were confirmed using denaturing gel electrophoresis to analyse the priming reaction products in the presence of increasing concentrations of Fludarabine-TP (Fig. 2A). We observed strong dose-dependent inhibition of RNA primer synthesis, with almost complete inhibition evident even at sub-stoichiometric concentrations of Fludarabine-TP compared to ATP. In the presence of an existing RNA primer annealed to a single-stranded DNA template, providing primase with Fludarabine-TP as the only available nucleotide limited primer extension to one or two nucleotides (Fig. 2B). Primase added the first Fludarabine moiety quickly (lane 4, first addition complete by 2 minutes), and the second much more slowly (lane 7, second addition complete by 30 minutes). An RNA primer with Fludarabine incorporated at its 3′-end could not be further extended with ATP, suggesting that incorporation and capping of the growing ribonucleotide chain is a likely mechanism of action (Fig. 2C). This is in agreement with previous primase studies which also describe chain termination by *ara* nucleotides (Huang et al., 1990; Catapano et al., 1993; Richardson et al., 2004).

**Figure 2:**
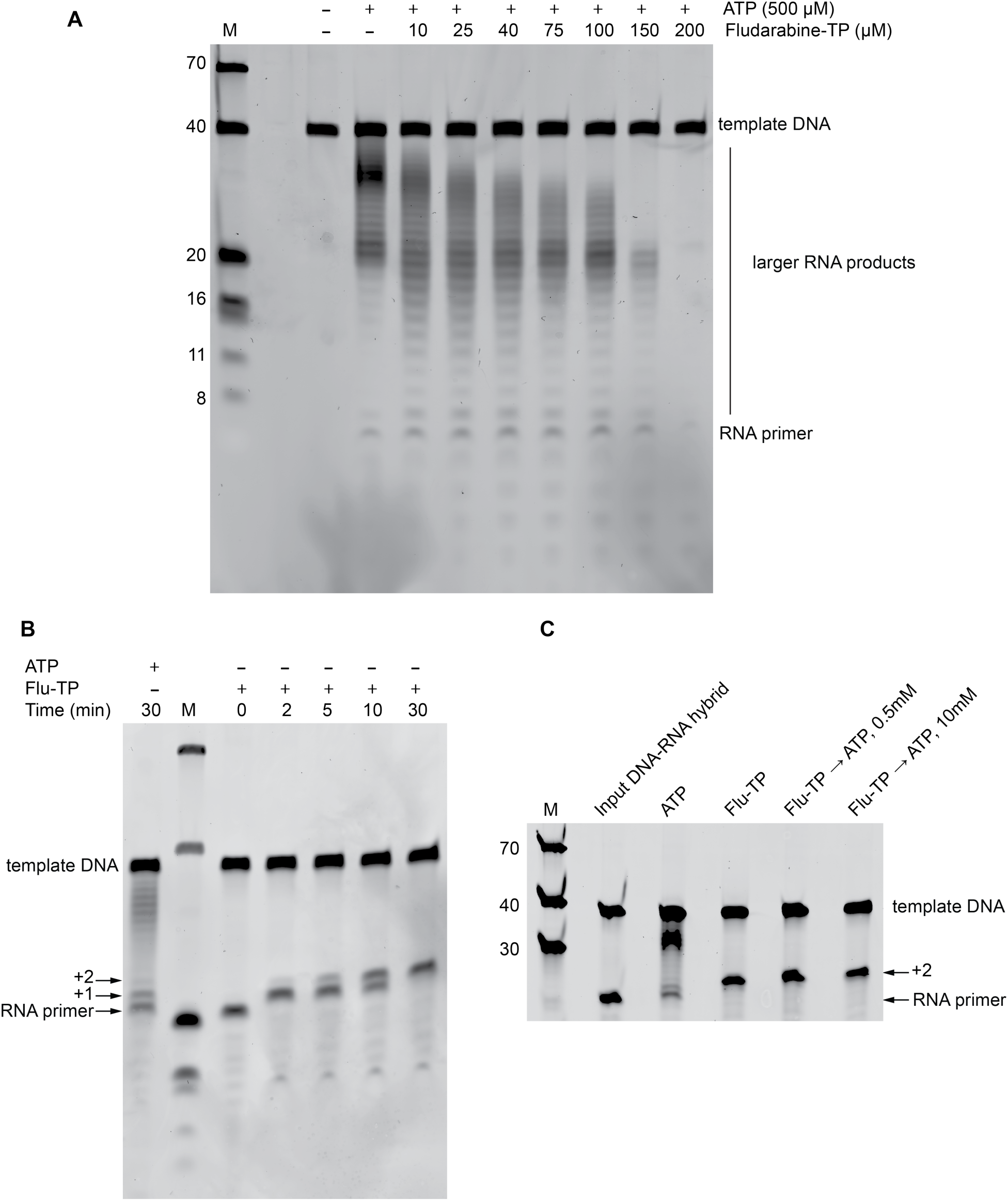
Effect of Fludarabine-TP on RNA primer synthesis and extension. (a) Denaturing gel showing the dose-dependent inhibition of RNA primer synthesis by Fludarabine-TP. Reactions contained 0.5 µM polydT40 ssDNA template, 0.5 µM primase, 500 µM ATP, 10mM Mg(OAc)_2_ and the indicated concentration of Fludarabine-TP. Reactions were incubated at 37 °C for 30 minutes, and the products were analysed by denaturing acrylamide gel electrophoresis. (b) Denaturing gel showing the incorporation of Fludarabine into an existing RNA primer. The template comprised a 38-mer DNA template (5′- T_20_CCAGAGAGCGCCCAAACG) annealed to an 18-mer RNA primer (5′- CGUUUGGGCGCUCUCUGG). Reactions contained 0.5 µM annealed DNA-RNA template, 0.5 µM primase, 10mM Mg(OAc)_2_, and 500 µM ATP or Fludarabine-TP. Reactions were incubated and analysed as in (a). (c) Denaturing gel showing that primase is unable to extend an RNA primer following Fludarabine incorporation. 0.5 µM primase was incubated with 0.5 µM DNA(38)-RNA(18) template and either 500 µM ATP (lane 3) or 500 µM Fludarabine-TP (lane 4) for 30 minutes at 37 °C. ATP (0.5 or 10 mM) was subsequently added to the Fludarabine-TP sample and incubated for a further 30 minutes (lanes 5,6). All gels were post-stained with Sybr Gold. M = marker.

We then used a fluorescence polarisation (FP) competition binding assay to compare the binding affinities of the various nucleotides and nucleotide analogues. The lower IC_50_ values obtained for the *ara* nucleotides (IC_50_^Fludarabine-TP^ = 1.13 μM, IC_50_^Vidarabine-TP^ = 1.45 μM) compared to those for the ribo/deoxyribonucleotides (IC_50_^ATP^ = 7.53 μM, IC_50_^2F-ATP^ = 7.19 μM, IC_50_^dATP^ = 3.34 μM), indicate that the *ara* nucleotides indeed bind with higher affinity (Figure 3A). Using thermal denaturation, we observed that all nucleotides stabilised primase relative to the unliganded enzyme (Fig. 3B). However, both Vidarabine-TP and Fludarabine-TP stabilised the primase to a greater extent relative to ATP (T_m_^ATP^ = 53.8 ± 0.3 °C, T_m_^Vidarabine-TP^ = 54.4 ± 0.2 °C, T_m_^Fludarabine-TP^ = 58.0 ± 0.2 °C). Surprisingly, binding of Fludarabine-TP resulted in significantly greater thermal stability than Vidarabine-TP, even though the competition experiments suggest very similar IC_50_ values.

**Figure 3:**
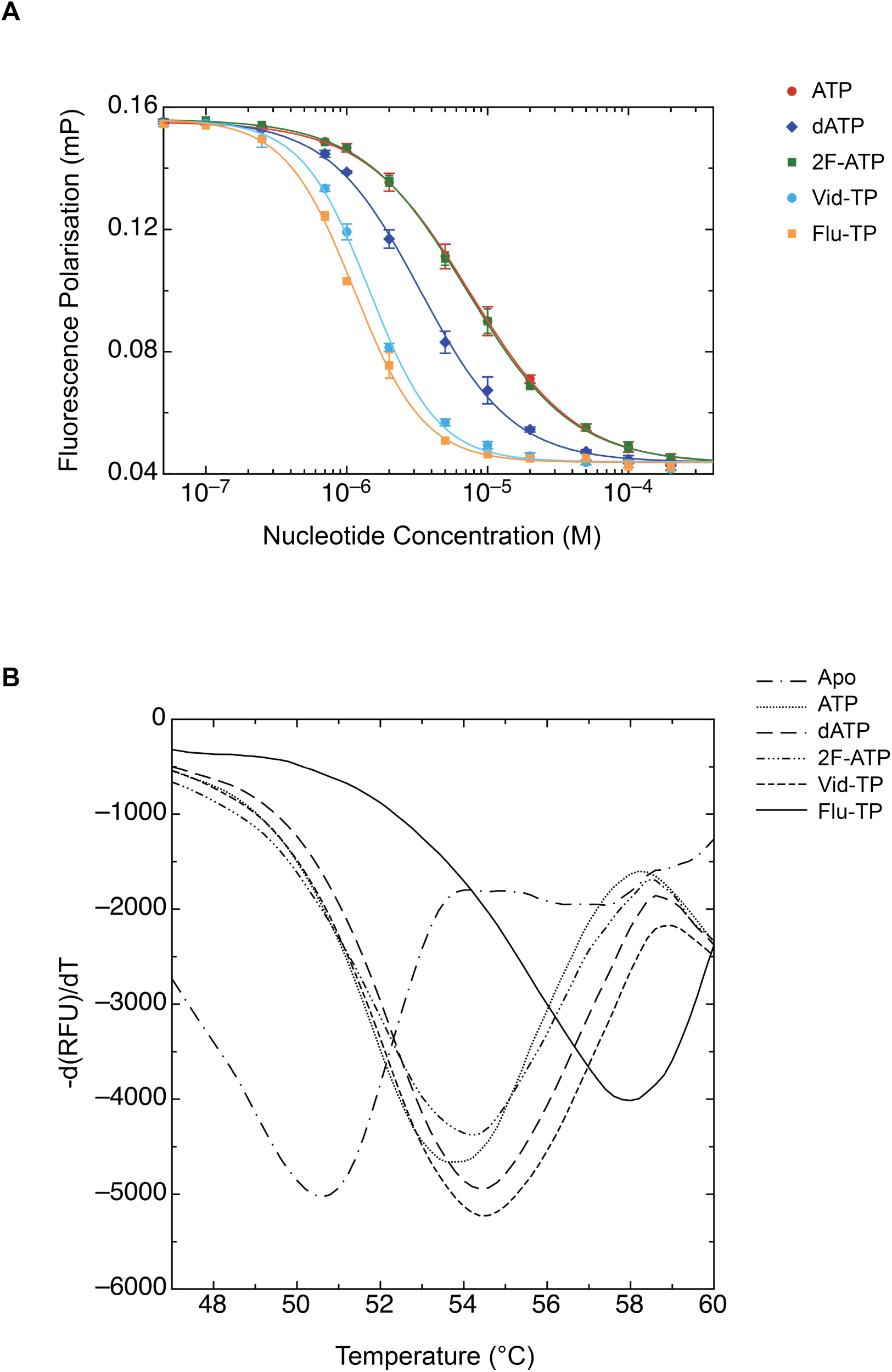
The *ara* nucleotides show enhanced binding to and stabilization of human primase. FP-based competition binding experiment in which Pri1 was pre-bound to 6FAM-labelled ATP and then challenged with increasing concentrations of the indicated nucleotide or nucleotide analogue. Each data point represents the mean ± s.d. (n=3). (b) First derivative of the thermal denaturation curve for the chimeric Pri1-Pri2^ΔCTD^-Pol a construct (see Methods) in the presence of the indicated nucleotide or nucleotide analogue. Single, representative curves are shown, although each melt was performed in quadruplicate. Calculated melting temperatures: T_m_^Apo^ (50.5 ± 0.2 °C), T_m_^ATP^ (53.8 ± 0.3 °C), T_m_^dATP^ (54.5 ± 0.1 °C), T_m_^2F-ATP^ (54.3 ± 0.1 °C), T_m_^Vid-TP^ (54.4 ± 0.2 °C), T_m_^Flu-TP^ (58.0 ± 0.2 °C) (mean ± s.d., n=4). RFU: relative fluorescence units.

### X-ray crystal structures of Fludarabine-TP and Vidarabine-TP bound to the Pri1 elongation site

Primase initiates RNA primer synthesis using two distinct nucleotide binding pockets: (i) the initiation site, which utilises several key residues on Pri2-CTD to bind the nucleotide that will be incorporated at the 5′-end of the primer and (ii) the Pri1 elongation site that binds all subsequent nucleotides (Baranovskiy et al., 2016; Copeland and Wang, 1993). Following synthesis of the first di-nucleotide, which involves both Pri1 and Pri2, further nucleotide addition requires only the elongation site on Pri1, (Klinge et al., 2007; Zerbe and Kuchta, 2002). To obtain atomic details of the interaction between *ara* nucleotides and the elongation site of human Pri1, we soaked Vidarabine-TP and Fludarabine-TP into Pri1 crystals that diffracted to near-atomic resolution and in which the active site was free from lattice contacts. For comparison, we determined in the same way the crystal structures of Pri1 bound to ATP, 2F-ATP and dATP. Given the higher affinity of primase for ATP in the presence of Mn^2+^ (K_d_^flu-ATP.Mn^ = 0.24 μM) compared to Mg^2+^ (K_d_^flu-ATP.Mg^ = 130 μM), we supplemented the nucleotide-containing crystal soak solutions with 500 μM MnCl_2_ (Supplementary Fig. 1). For each nucleotide soak, the electron density map clearly revealed a single nucleotide bound to the elongation pocket of Pri1 (Supplementary Fig. 2 A-E). Data collection and refinement statistics are given in Table 1.

**Table 1:** Data collection and refinement statistics for the x-ray crystal structures of Pri1 bound to nucleotides and nucleotide analogues

|  | **ATP** | **dATP** | **2F-ATP** | **Vid-TP** | **Flu-TP** |
| --- | --- | --- | --- | --- | --- |
| Accession code | 6R4S | 6R5D | 6R5E | 6R4T | 6R4U |
| **Data collection** |  |  |  |  |  |
| Wavelength (Å) | 0.979 | 0.917 | 1.039 | 0.979 | 0.979 |
| Space group | C222<sub>1</sub> | C222<sub>1</sub> | C222<sub>1</sub> | C222<sub>1</sub> | C222<sub>1</sub> |
| **Cell dimensions** |  |  |  |  |  |
| a, b, c (Å) | 110.1<br>117.3<br>151.2 | 110.5<br>117.3<br>151.7 | 110.5<br>117.2<br>152.1 | 111.1<br>119.0<br>150.7 | 110.8<br>117.6<br>148.8 |
| $\alpha, \beta, \gamma$ (°) | 90, 90, 90 | 90, 90, 90 | 90, 90, 90 | 90, 90, 90 | 90, 90, 90 |
| Molecules/asymmetric unit | 2 | 2 | 2 | 2 | 2 |
| Resolution (Å) | 29.32-2.75<br>(2.91-2.75) | 46.40-1.95<br>(2.07-1.95) | 46.41-1.85<br>(1.96-1.85) | 46.69-2.35<br>(2.49-2.35) | 46.13-2.20<br>(2.33-2.20) |
| Unique reflections | 25616<br>(2462) | 71418<br>(11299) | 82430<br>(12971) | 41814<br>(6609) | 49380<br>(7541) |
| R<sub>meas</sub> | 0.18 (0.92) | 0.09 (1.20) | 0.08 (0.77) | 0.08 (1.05) | 0.10 (1.38) |
| Mean I/σI | 5.46<br>(1.04) | 14.56<br>(1.33) | 11.52<br>(1.52) | 15.35<br>(1.63) | 12.91<br>(1.52) |
| Completeness, % | 96.8 (94.5) | 99.2 (98.0) | 97.7 (95.8) | 99.7 (98.7) | 99.2 (95.3) |
| Redundancy | 4.95 | 6.75 | 4.39 | 7.20 | 6.50 |
| CC<sub>1/2</sub> | 0.98 (0.36) | 1.00 (0.75) | 1.00 (0.77) | 1.00 (0.81) | 1.00 (0.83) |
| Wilson B-factor | 67.09 | 37.03 | 34.23 | 62.01 | 44.73 |
| **Refinement** |  |  |  |  |  |
| Non-hydrogen atoms | 6543 | 7057 | 7028 | 6642 | 6738 |
| R<sub>work</sub>/R<sub>free</sub>, % | 22.7/26.8<br>(35.7/36.1) | 17.6/21.5<br>(40.1/41.9) | 17.5/20.5<br>(37.4/37.6) | 20.8/23.8<br>(42.0/50.8) | 19.4/22.3<br>(59.0/65.5) |
| Average B factor | 71.08 | 50.37 | 46.40 | 79.78 | 59.06 |
| Clashscore | 2.87 | 1.36 | 1.59 | 1.84 | 1.69 |
| **Rmsd** |  |  |  |  |  |
| Bond lengths | 0.004 | 0.003 | 0.003 | 0.003 | 0.004 |
| Bond angles | 0.81 | 0.57 | 0.68 | 0.62 | 0.58 |
| **Ramachandran analysis** |  |  |  |  |  |
| Preferred region, % | 96.70 | 96.98 | 97.09 | 96.98 | 97.35 |
| Allowed regions, % | 3.30 | 2.89 | 2.91 | 3.02 | 2.65 |
| Outliers, % | 0.00 | 0.13 | 0.00 | 0.00 | 0.00 |

Statistics in parentheses indicate those for the highest resolution shell.

This high-resolution structural information allowed us to compare in detail the interactions of the different sugar moieties within the active site. As described previously, human Pri1 adopts the mixed α/β primase (Prim) fold characteristic of eukaryotic and archaeal primases, with a catalytic triad comprising Asp109, Asp111 and Asp306 which together co-ordinate two divalent metal ions (Kilkenny et al., 2013; Copeland and Tan, 1995). As seen in previous nucleotide-bound Pri1 crystal structures, the triphosphate moiety of the nucleotide resides in a basic pocket formed by the side-chains of residues Arg162, Arg163, His166, Lys318 and His324, as well as the two Mn^2+^ ions co-ordinated by the catalytic aspartates (Supplementary Fig. 2F) (Kilkenny et al., 2013; Vaithiyalingam et al., 2014). By superposing the apo and nucleotide-bound structures, we observe that the loop containing Asp306 needs to move towards the active site to allow the Asp306 side-chain to co-ordinate the second Mn^2+^ ion effectively (Supplementary Fig. 3). Interestingly, while the ATP, dATP, 2F-ATP and Fludarabine-TP structures show a fully engaged loop, the Vidarabine-TP structure reveals the loop to adopt an intermediate or apo conformation, and the second Mn^2+^ ion could not be reliably modeled for either chain of the asymmetric unit (Supplementary Fig. 2D).

In all structures, the sugar moiety is positioned by the formation of a hydrogen bond between the ribose 3′-OH and the main-chain amide NH of Lys318 (Fig. 4A). This interaction is important for catalysis because Cordycepin-TP (3′- deoxyadenosine triphosphate) is not efficiently polymerised onto an existing RNA primer and only starts to inhibit RNA primer synthesis when present in excess of ATP (Supplementary Fig. 4). In addition to this hydrogen bond, C4 and C5 of the ribose moiety pack on top of the aliphatic side-chain of Leu317. In the ATP- and 2F-ATP-bound structures, the 2′-OH of the ribose inserts between the main-chain carbonyl group of Leu316 and the carboxyl group of the Asp79 side-chain and is positioned roughly equidistant between these two moieties. However, in the Vidarabine-TP and Fludarabine-TP bound structures, the 2′-OH points directly towards the carboxyl group of Asp79 and the ε-amino group of Lys77, and away from the carbonyl O of Leu316 (Fig. 4A). In this way, the *ara* nucleotides present the 2′-OH in an orientation that is more favourable for hydrogen bond formation compared to the normal ribonucleotides (distances from *ara* 2′-OH to Asp79 carboxylate-O range from 2.9 to 3.3 Å and from *ara* 2′-OH to Lys77 ε-N range from 2.8 to 3.5 Å). The fluorine atom on the base moiety of Fludarabine-TP resides above the side-chain of Leu316 (Fig. 4A), and only 3.6-3.9 Å away from the side-chain of the catalytically essential residue Arg56 (Kilkenny et al., 2013). This pocket could certainly be explored further in future structure-based drug design efforts.

**Figure 4:**
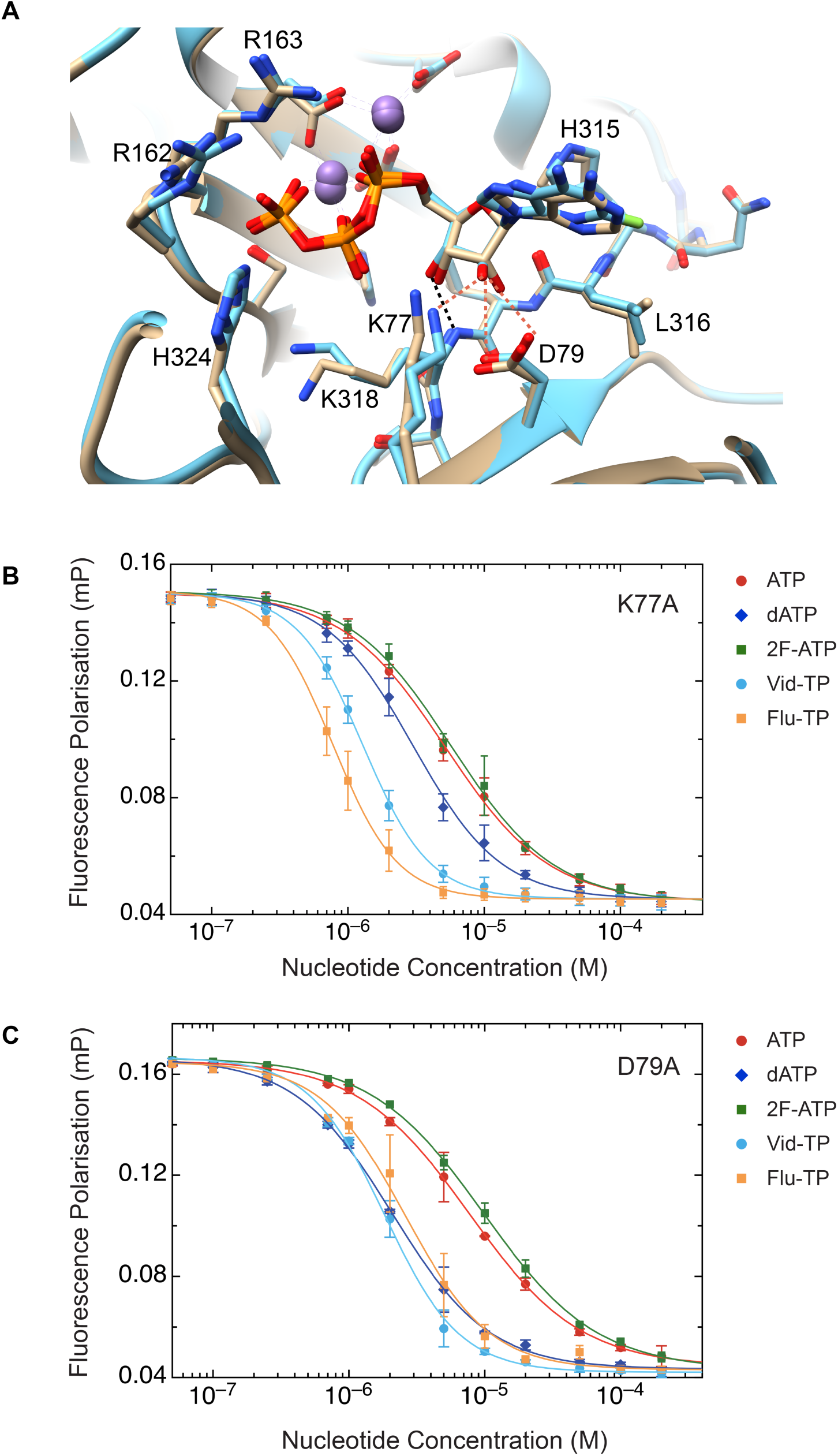
Asp79 contributes to the preference of primase for binding *ara* nucleotides. (a) Superposition of the x-ray crystal structures of Pri1 bound to ATP (beige) and Fludarabine-TP (blue). In both structures the ribose 3′-OH forms a hydrogen bond to the main chain NH group of Lys318 (dashed line, black). In the Fludarabine-TP structure the ribose 2′-OH is poised to interact with the side-chains of Asp79 and/or Lys77 (dashed lines, orange). Mn^2+^ ions are shown as purple spheres. Image generated using Chimera (Pettersen et al., 2004). FP-based competition binding experiment in which Pri1 point mutant K77A was pre-bound to 6FAM-labelled ATP then challenged with increasing concentrations of the indicated nucleotide or nucleotide analogue. Each data point represents the mean ± s.d. (n=3). (c) As in (b) but with Pri1 point mutant D79A.

The Vidarabine-TP and Fludarabine-TP crystal structures indicate that the side-chain of residues Lys77 and/or Asp79 may be responsible for the observed preference of primase for *ara* nucleotides. To test this hypothesis, we generated and purified K77A and D79A point mutants of Pri1. Size exclusion chromatography was used to confirm that these mutants were not mis-folded (Supplementary Fig. 5). We then performed nucleotide binding experiments with the Pri1 point mutants, using the FP competition assay described above. The K77A mutant showed comparable IC_50_ curve profiles to wild-type Pri1 (Fig. 4B). In the case of the D79A mutant, however, the competition curves for the *ara* nucleotides were indistinguishable from that of dATP, with the IC_50_ for Fludarabine-TP increasing from 1.13 μM to 2.60 μM, and the IC_50_ for dATP decreasing from 3.34 μM to 2.09 μM (Fig. 4C). These results indicate that Asp79 does play a role in distinguishing between different configurations of the sugar moiety. However, the D79A point mutant was still inhibited by Fludarabine-TP and Vidarabine-TP (Supplementary Fig. 6C), possibly indicating some redundancy between Asp79 and Lys77 in providing hydrogen bonding to the *ara* 2’-OH of the ribose.

### The ara 2′-OH is crucial for mediating effective primase inhibition

To investigate whether primase inhibition is a shared property of chemotherapeutic nucleotide analogues, we examined RNA primer synthesis activity in the presence of an extended repertoire of these agents, including Gemcitabine-TP, Cytarabine-TP and Clofarabine-TP (Fig 1A). On a ssDNA template, titration of Cytarabine-TP resulted in very similar levels of primase inhibition to Fludarabine-TP (Fig. 5A). Given that Cytarabine-TP and Fludarabine-TP are CTP and ATP analogues, respectively, this experiment was conducted using a ssDNA template containing equal quantities of T and G. This result was corroborated in a fluorescence-based primer synthesis assay, which showed similar levels of inhibition for Fludarabine-TP, Vidarabine-TP and Cytarabine-TP (Fig. 5B). In addition, Cytarabine was readily incorporated into an existing RNA primer, as described previously (Supplementary Fig. 7B) (Richardson et al., 2004). These results further confirm that primase inhibition by *ara* 2′-OH nucleotide analogues is largely unaffected by the nature of the base moiety.

**Figure 5:**
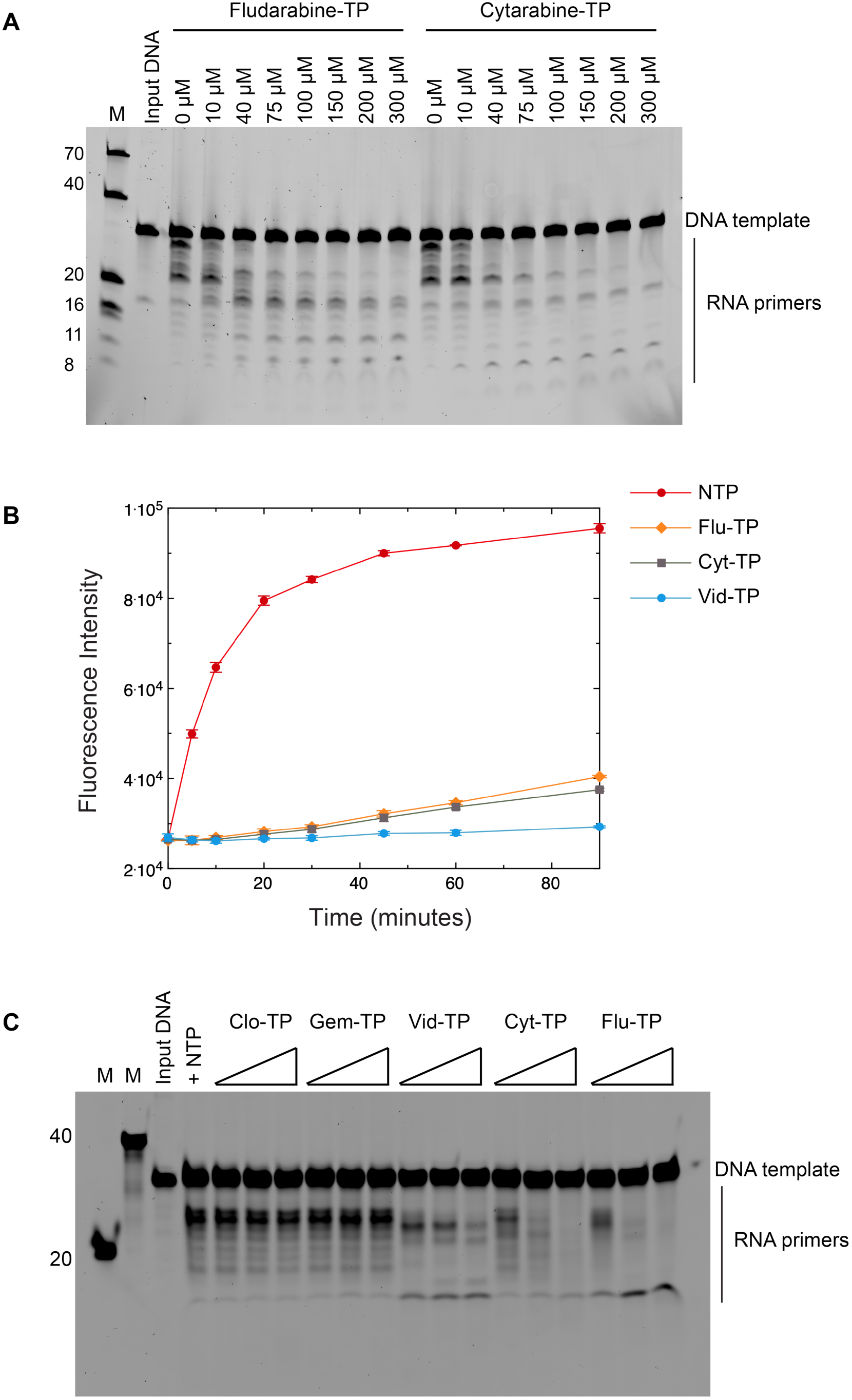
*ara* 2’-OH but not *ara* 2′-F nucleotide analogues mediate efficient inhibition of RNA primer synthesis. (a) Denaturing gel showing the effect of Fludarabine-TP and Cytarabine-TP on RNA primer synthesis. Reactions contained 0.5 µM primase, 0.5 µM ssDNA template (5′- ATGAGTGAATGTCTGTGAGTGTCTGCCTGC), 500 µM each NTP, 10 mM Mg(OAc)_2_ and the indicated concentration of nucleotide analogue. Reactions were incubated at 37 °C for 30 minutes then analysed by denaturing polyacrylamide gel, post-stained with Sybr Gold. (b) Fluorescence-based RNA primer synthesis assay on the same ssDNA template described in (a) above, in the presence of Mn^2+^ ions and equimolar concentrations of ribonucleotides (20 µM each NTP) and nucleotide analogue (20 µM): Fludarabine-TP (Flu-TP), Cytarabine-TP (Cyt-TP) or Vidarabine-TP (Vid-TP). Each data point represents the mean ± s.d. (n=3). (c) Denaturing gel showing the effect of the 2′-F *ara* nucleotide analogues, Clofarabine-TP (Clo-TP) and Gemcitabine-TP (Gem-TP), on RNA primer synthesis. Reactions contained 1 µM ssDNA template (5′- GTTGTCCATTATGTCCTACCTCGTGCTCCT), 0.5 µM primase, 500 µM each NTP, 10mM Mg(OAc)_2_ and the indicated nucleotide analogue (25, 100 or 500 µM). Reactions were incubated and analysed as in (a). M = marker.

By contrast, Gemcitabine-TP and Clofarabine-TP, which both contain an *ara* 2′-F rather than *ara* 2′-OH, did not inhibit RNA primer synthesis over the range of concentrations tested (Fig. 5C). In addition, in an FP-based competition binding experiment, Gemcitabine-TP and Clofarabine-TP showed very similar IC_50_ values to ATP (Supplementary Fig. 7 A). This is consistent with the structural data presented here, as an *ara* 2′-F would be unable to form a favourable hydrogen bond with the carboxylate side-chain of Asp79. Interestingly, in the presence of nucleotide analogue alone, while Clofarabine was poorly incorporated into an existing RNA primer, Gemcitabine was readily incorporated (Supplementary Fig. 7B). We conclude that, while primase can incorporate Gemcitabine into RNA primers, this analogue does not exert significant inhibition of primer synthesis in the presence of the natural ribonucleotides. This is probably due to its weaker binding affinity compared to the *ara* 2′-OH nucleotide analogues. Taken together, these results indicate that a hydrogen bond donor in the 2′ position of the arabinose sugar moiety is crucial for mediating effective primase inhibition by these nucleotide analogues.

## DISCUSSION

We have analysed in detail the mode of binding of anti-cancer agent Fludarabine-TP and anti-viral agent Vidarabine-TP to human primase. Thermal denaturation, competition and primer synthesis experiments all confirm that these arabinofuranosyl nucleotides are *bona fide* inhibitors of primase, as is the CTP- analogue, Cytarabine-TP. We find that Asp79 contributes to the preference of primase to bind *ara* nucleotides over ribo- and deoxyribonucleotides, and attribute this, at least in part, to the formation of a favourable interaction between the *ara* 2′-OH and the carboxylate side-chain of Asp79 (Fig. 4A).

Given that these *ara* nucleotides inhibit primase to a significant degree in the presence of excess normal ribonucleotides *in vitro* (Fig. 2A, Fig. 5A), and the fact that Fludarabine-TP can accumulate to concentrations in the range of hundreds of micromolar inside cells (Gandhi and Plunkett, 2002), it is possible that primase may indeed be one of the relevant targets of this chemotherapeutic agent *in vivo*. Whether primase inhibition by Fludarabine-TP or Cytarabine-TP is one of the primary modes of cytotoxicity towards cancer cells remains to be determined, and will be the subject of future experiments.

Gemcitabine, Cytarabine, Fludarabine and Clofarabine are all used as cytotoxic anti-cancer agents in the clinic but, despite structural similarities, their mode of action seems to vary quite significantly. A recent structural and biochemical study of the interaction of various chemotherapeutic nucleotide analogues with SamHD1, an enzyme that regulates dNTP levels, has important implications for drug turnover and hence efficacy (Srikanth et al., 2018). In addition, human PrimPol has been shown to incorporate Cytarabine-TP and Gemcitabine-TP, but not two antiviral nucleoside analogues (Emtricitabine and Lamivudine), into newly-synthesised DNA strands (Tokarsky et al., 2017). These recent studies highlight how important it will be to analyse the effect of each nucleotide analogue on a range of intracellular targets in order to begin to understand their nuanced activity *in vivo*.

The data presented here indicate that chemotherapeutic nucleotides Cytarabine-TP and Fludarabine-TP, but not Gemcitabine-TP or Clofarabine-TP, are likely to be effective inhibitors of primase activity *in vivo*. They display higher binding affinity to human primase than the natural ribonucleotides (Fig. 3A), are efficiently incorporated into RNA primers (Fig. 2B, Supplementary Fig. 7B) and have the potential to induce chain termination of nascent RNA primers (Fig. 2C). Thus it is likely that their inhibitory action is a result of both competitive inhibition and capping of the RNA primer resulting in chain termination.

It should now be possible to build on previous work outlining the synthesis of novel *ara* nucleotide analogue compounds as potent primase inhibitors (Moore et al., 2002). Initial work was promising, describing a nucleotide analogue (araBTP) with improved selectivity for primase over DNA polymerase a, but further advances were hindered by a lack of structural information relating to the mode of binding of this agent to the primase active site. Based on the structural data presented here, we hypothesise that it should be possible to further optimise the binding of *ara* nucleotides to Pri1, in order to generate a higher affinity, more selective inhibitor of human primase. In the longer term, this may result in a new chemotherapeutic agent that has less severe side-effects than Fludarabine, by minimising cytotoxicity that results from off-target effects. In addition, structural information of this kind will enable the refinement of individual therapeutic nucleotide analogues so that they interact with specific combinations of intracellular targets, thereby modulating their effect *in vivo*.

## MATERIALS AND METHODS

### Nucleotides and nucleotide analogues

2F-ATP (NU-145S), Vidarabine-TP (NU-1111S), Fludarabine-TP (NU-10703-10), Gemcitabine-TP (NU-1607S), Clofarabine-TP (NU-874), Cytarabine-TP (NU- 1170S) and N^6^-(6-Aminohexyl)-ATP-6FAM (NU-805-6FM) were purchased from Jena Bioscience.

### Cloning, expression and protein purification

Heterodimeric human primase (Pri1-Pri2) was co-expressed in bacteria as full-length His_6_-tagged Pri1 (amino acids 1 to 420) and Pri2 (1 to 462). Residues 463 to 509 of Pri2 were omitted to minimise proteolytic degradation. These residues are not conserved and are disordered in the x-ray crystal structure of full-length human primase (Baranovskiy et al., 2016). Primer synthesis assays confirmed that this protein showed similar activity to the wild-type protein (Supplementary Fig. 8). Human Pri1 was produced as either full-length His_10_-tagged Pri1 (1 to 420) for competition experiments, or His_10_-tagged Pri1 (1 to 407) for crystallisation. K77A and D79A point mutations in full-length Pri1 were generated by site-directed mutagenesis. Due to exposed hydrophobic patches on both Pri1 and Pri2 that interfered with the analysis, thermal melt experiments were performed using a chimeric construct of human primase (Pri1-Pri2^ΔCTD^-Pol a chimera), comprising Pol a residues 1445 to 1462 (GYSEVNLSKLFAGCAVKS) fused via a 15 residue Gly-Ser-Thr linker to residue N19 of Pri2. In this protein, the tethered Pol a peptide binds to a hydrophobic patch on the N-terminal domain of Pri2, while the Pri1 elongation site is unaffected (Kilkenny et al., 2013). All proteins were expressed in the Rosetta2 (DE3) *E. coli* strain from the pRSFDuet-1 vector (Novagen). The purification protocol entailed Ni-NTA agarose chromatography (Qiagen), Heparin sepharose chromatography (GE Healthcare), His-tag cleavage by TEV protease, and size exclusion chromatography. The size exclusion buffer comprised either 25 mM HEPES pH 7.2, 300 mM KCl, 5% glycerol and 1 mM TCEP (for Pri1-Pri2), or 25 mM HEPES pH 7.2, 150 mM KCl, 5% glycerol and 1 mM TCEP (for Pri1). Purified proteins were concentrated and aliquots flash frozen in liquid nitrogen, then stored at −80 °C.

### Crystallisation and X-ray crystallography

Apo Pri1 (1 to 407) was crystallised by vapour diffusion at 19 °C, by mixing 1 μl of 150 μM Pri1 with 1 μl of crystallisation buffer (0.1 M Bis Tris propane pH 6.5, 24% PEG 3350, 0.15 M NaF). Diffraction data were collected at SOLEIL synchrotron (beamline PROXIMA 1). The protein crystallized in space group P4_3_2_1_2, with one copy of Pri1 in the asymmetric unit. Diffraction data were indexed, integrated and scaled using XDS (Kabsch, 2010), and the structure solved by molecular replacement in Phaser (McCoy et al., 2007) using 4BPU as the search model (Kilkenny et al., 2013). The model was completed by alternating between cycles of manual rebuilding in Coot (Emsley et al., 2010) and structure refinement in PhenixRefine (Adams et al., 2010). Unfortunately, while these crystals diffracted to high resolution (1.5 Å), they were often multiple and/or poorly reproducible, and the crystallisation condition was therefore optimised for the nucleotide soaking experiments. Pri1 was subsequently crystallised by vapour diffusion at 19 °C, by mixing 1.5 μl of 150 μM Pri1 with 1 μl of crystallisation buffer (23% PEG 3350, 10% ethylene glycol, 200 mM Na/K tartrate). Crystals were soaked overnight in crystallisation buffer containing 500 μM nucleotide (ATP, dATP, 2F-ATP, Vidarabine-TP or Fludarabine-TP) and 500 μM MnCl_2_. Diffraction data were collected at beamlines I24 (ATP, 2F-ATP and Vidarabine-TP), I04-1 (dATP) and I02 (Fludarabine-TP) of the Diamond Light Source, UK. The protein crystallized in space group C222_1_, with two copies of Pri1 in the asymmetric unit (Table 1). The apo Pri1 structure described above was used as the molecular replacement model. Data were processed and the model refined as described above. Owing to poor electron density, the following residues were considered disordered and omitted from the final model: apo Pri1 (chain A: 360-381); Pri1.ATP (chain A: 1-2, 360-381, 407; chain E: 1, 360-381, 407), Pri1.dATP (chain A: 1-4, 361-379; chain E: 1, 361-381), Pri1.2F-ATP (chain A: 1-5, 361-379; chain E: 1-3, 361-381, 407), Pri1.Vidarabine-TP (chain A: 1-5, 362-379; chain D: 1, 361-381), Pri1.Fludarabine-TP (chain A: 1-5, 361-380; chain E: 1-3, 360-381).

### Fluorescence-based primase activity assays

Time-course RNA primer synthesis assays were performed in triplicate in 96 well plate format. Each well contained 1 μM Pri1-Pri2 (WT or D79A) and 200 nM ssDNA (5′-GTTGTCCATTATGTCCTACCTCGTGCTCCT) in 25 mM HEPES pH 7.0, 120 mM NaCl, 1 mM TCEP, 5 mM MgCl_2_ (or 1mM MnCl_2_). Reactions were initiated by the addition of 20 μM of each ribonucleotide (ATP, CTP, GTP and UTP) and 20 μM nucleotide analogue (ATP, dATP, 2F-ATP, Vidarabine-TP or Fludarabine-TP), to give a final reaction volume of 25 μl. The plate was incubated at 37 °C in the presence of MgCl_2_, or 25 °C in the presence of MnCl_2_. Reactions were quenched at the indicated time points by the addition of 25 μl 1:100 dilution of PicoGreen (Thermo Fisher Scientific) in 25 mM Tris pH 8.0 and 20 mM EDTA. Following incubation of the plate at 25 °C for 10 minutes, fluorescence intensity measurements were recorded in a PHERAstar Plus multi-detection plate reader (BMG Labtech) equipped with fluorescence intensity optic module (λ_ex_ = 485 nm; λ_em_ = 520 nm). Each data point is the mean of 20 flashes per well. The voltage gain was set by adjusting the fluorescence intensity of a well containing protein, nucleotide and 30-mer dsDNA (ssDNA as above, with annealed complementary strand), to 90% of the maximum measurable intensity.

### Gel-based primase activity assays

Each 30 μl reaction contained 20 mM Tris-HCl pH 7.5, 50 mM K(OAc), 1 mM DTT, 0.5 mM NTP, 0.5 μM ssDNA or annealed DNA-RNA template, 0.5 μM Pri1-Pri2, the indicated concentration of nucleotide analogue and either 10 mM Mg(OAc)_2_ or 1 mM MnCl_2_. The control reaction contained all components except primase. Reactions were incubated at 37 °C for the indicated time to allow primer synthesis to occur, then terminated by the addition of 30 μl buffer comprising 95% formamide and 25 mM EDTA. Samples were heated to 70 °C for 2 minutes then loaded onto a 18% urea-polyacrylamide gel, which was run at 500 V for 90 minutes in 0.5x TBE buffer. Gels were stained in 0.5x TBE buffer containing a 1:10000 dilution of Sybr Gold Stain (Thermo Fisher Scientific), for 30 min with shaking. Reaction products were visualised by scanning with a 473 nm laser (Typhoon FLA 9000, GE Healthcare).

### Fluorescence anisotropy-based nucleotide binding experiments

Binding experiments were performed in triplicate in 96 well plate format. Each well contained 30 nM 6FAM-labelled ATP in 25 mM HEPES pH 7.0, 120 mM NaCl, 1 mM TCEP and 2 mM MnCl_2_ (or 2 mM MgCl_2_). Pri1-Pri2^ΔCTD^-Pol a chimera was added in increasing concentrations, ranging from 0 to 120 μM (in the presence of MgCl_2_), or 0 to 5.8 μM (in the presence of MnCl_2_). Fluorescence anisotropy measurements were recorded in a PHERAstar Plus multi-detection plate reader (BMG Labtech) equipped with fluorescence polarization optic module (λ_ex_ = 485 nm; λ_em_ = 520 nm), at 25 °C. Each data point is the mean of 200 flashes per well. The voltage gain was set by adjusting the target mP values of fluorescein-labelled ATP relative to that of fluorescein (35 mP).

### Fluorescence anisotropy-based competition binding experiments

Competition binding experiments were performed in triplicate in 96 well plate format. Each well contained 30 nM 6FAM-labelled ATP and 1.5 μM Pri1, and nucleotide analogue was titrated in in increasing concentrations, from 0 to 200 μM. Binding buffer comprised 25 mM HEPES pH 7.0, 120 mM NaCl, 1 mM TCEP and 1 mM MnCl_2_. Fluorescence anisotropy measurements were recorded in a PHERAstar Plus multi-detection plate reader (BMG Labtech), as described above.

### Thermal denaturation

Reactions were performed in quadruplicate, in 96 well plate format. Each well contained 6.3 μM Pri1-Pri2^ΔCTD^-Pol a chimera, 500 μM MnCl_2_, 500 μM nucleotide and 5x SYPRO orange dye (Thermo Fisher Scientific), in a reaction buffer comprising 25 mM HEPES pH 7.0, 150 mM NaCl and 1 mM TCEP. Heating was performed in a CFX Connect Real-time PCR detection system with 96-well reaction module (Bio-Rad), using a ramp rate of 0.2 °C.min^-1^. The negative of the first derivative of the fluorescence signal was plotted against temperature, and the melting temperature (T_m_) estimated from the local minimum of the curve peak.

### Size exclusion chromatography

Protein samples were injected onto a Superdex S200 or S75 16/60 size exclusion column (GE Healthcare), which was run at 1 ml.min^-1^ in 25 mM HEPES pH 7.0, 200 mM NaCl and 5% glycerol. Eluted fractions were analysed by SDS-PAGE with Coomassie staining.

## ACCESSION NUMBERS

The co-ordinates and structure factors for the x-ray crystal structures presented in this paper have been deposited in the PDB under accession codes 6R4S (Pri1.ATP), 6R5D (Pri1.dATP), 6R5E (Pri1.2F-ATP), 6R4T (Pri1.Vidarabine-TP), 6R4U (Pri1.Fludarabine-TP), 6RB4 (apo Pri1).

## ACKNOWLEDGEMENTS

We thank Diamond Light Source for access to beamlines I02, I04-1 and I24 (MX9537) that contributed to the results presented here. We also acknowledge SOLEIL for provision of synchrotron radiation facilities and would like to thank the staff of beamline PROXIMA1 for assistance. We are grateful to the Facility Managers of the Crystallographic X-ray Facility (Dr Dimitri Chirgadze) and the Biophysics Facility (Dr Katherine Stott) at the Department of Biochemistry, University of Cambridge for their assistance in using these facilities. We are grateful to Dr Joseph Maman for expert advice on all fluorescent measurements. This work was funded by a Wellcome Trust Investigator Award (104641/Z/14/Z) to LP. SH was funded by a Boehringer-Ingelheim Fonds PhD Fellowship, Janggen-Pöhn-Stiftung Award and Swiss National Science Foundation Award.

## AUTHOR CONTRIBUTIONS

MLK designed the experiments and analysed results. MLK performed the fluorescence-based primer synthesis assays. MLK and SH performed the competition binding experiments. MLK and IRS performed the Cytarabine-TP, Gemcitabine-TP and Clofarabine-TP primer synthesis assays. MJB optimised the gel-based primer synthesis assay and carried out the initial Fludarabine-TP and Vidarabine-TP studies. MLK and NJR collected and processed the x-ray diffraction data. LP and MLK conceived the project and directed the research. MLK wrote the manuscript, with contributions from all authors.

