## Supplementary Figures for "Structural basis for inhibition of human primase by arabinofuranosyl nucleoside analogues Fludarabine and Vidarabine"

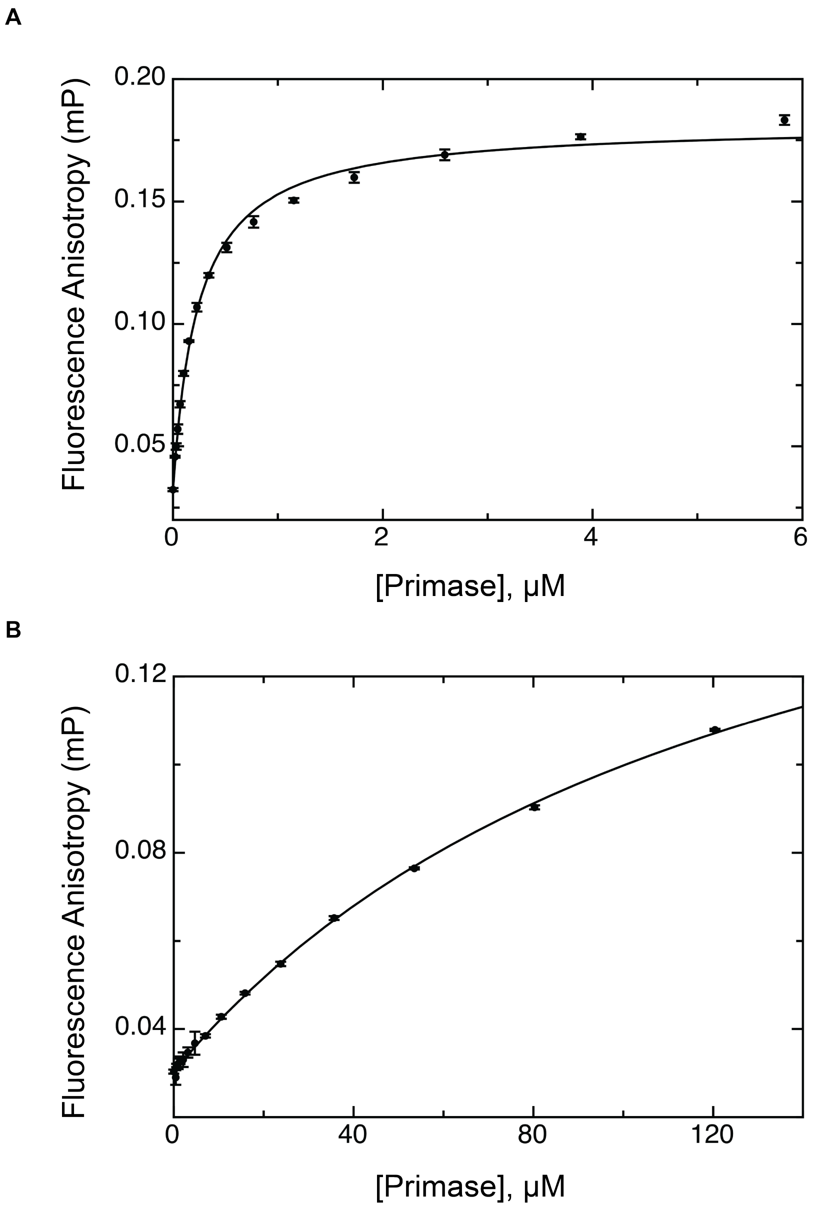


**Supplementary Figure 1**: $Mn^{2+}$ stimulates nucleotide binding to primase

Fluorescence polarisation was used to determine the binding affinity of primase for 6FAM-ATP in the presence of (a) $Mn^{2+}$ ions or (b) $Mg^{2+}$ ions. Each data point represents the mean ± s.d. (n=3).

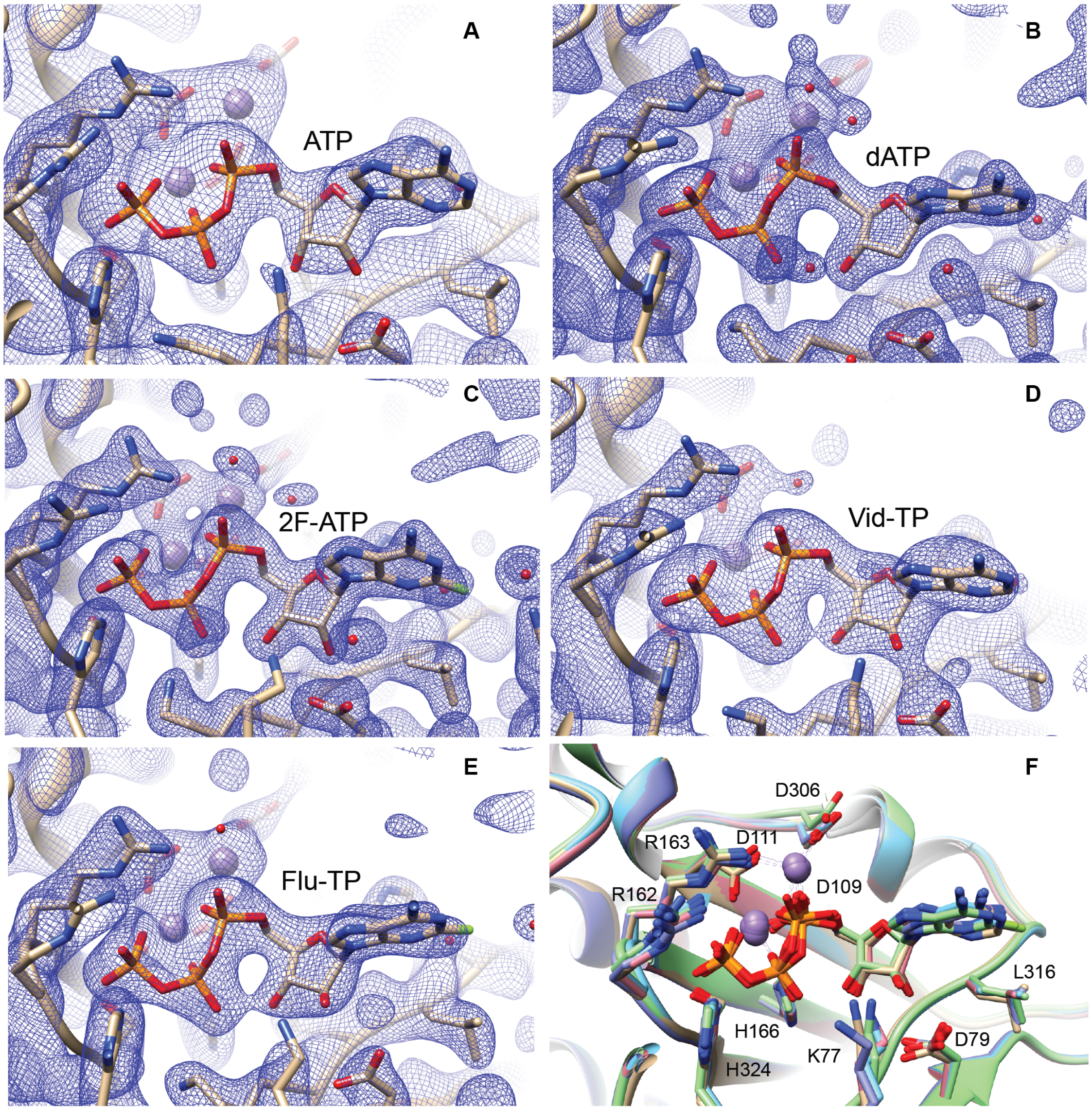


**Supplementary Figure 2**: Electron density maps showing nucleotides and nucleotide analogues bound to the elongation site of Pri1. 2Fo-Fc electron density map (contoured at 1.0 σ) showing the elongation site of Pri1 in complex with $Mn^{2+}$ ions (grey spheres) and (a) ATP, (b) dATP, (c) 2F-ATP, (d) Vidarabine-TP and (e) Fludarabine-TP. (f) Superposition of the x-ray crystal structures of Pri1 bound to ATP (beige), dATP (purple), 2F-ATP (pink), Vidarabine-ATP (green) and Fludarabine-TP (blue). Images generated using Chimera (Pettersen et al., 2004).

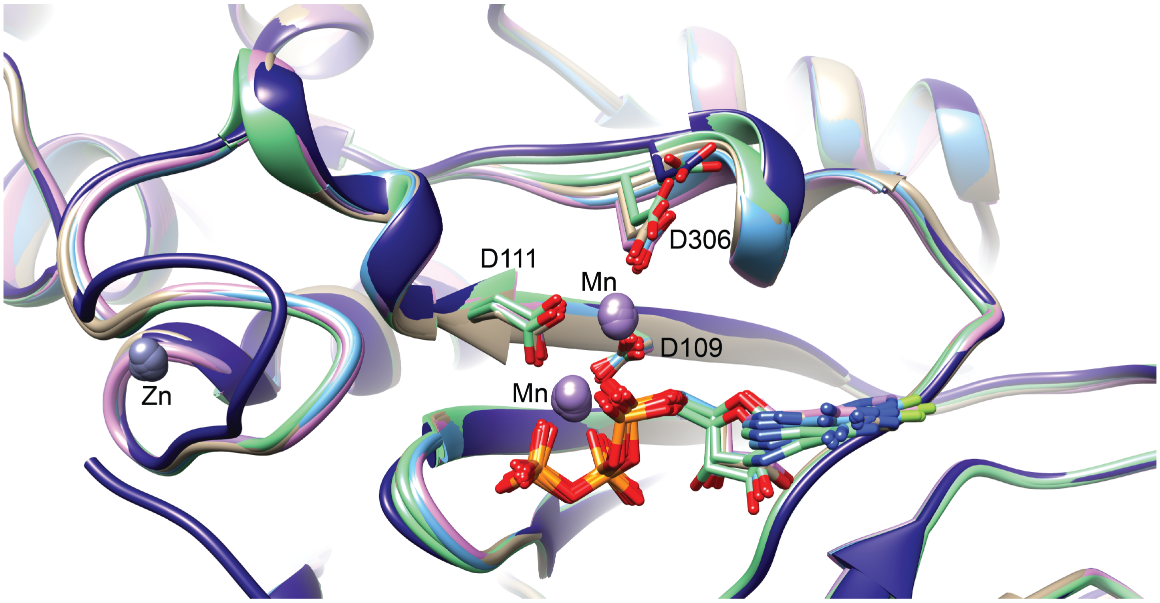


**Supplementary figure 3**: Superposition of the x-ray crystal structures of apo and nucleotide-bound Pri1. Superposition shows the elongation site of Pri1 in complex with and ATP (beige), dATP (purple), 2F-ATP (pink), Vidarabine-ATP (green) or Fludarabine-TP (light blue). Both Pri1 chains in the asymmetric unit of the nucleotide-bound structures are shown. $Mn^{2+}$ ions are shown as purple spheres. The single chain of the apo structure is shown in dark blue. Conformational changes in the loop co-ordinating the $Zn^{2+}$ ion are most likely due to differences in crystal packing between the apo and nucleotide-bound structures. Co-ordinated $Zn^{2+}$ ions are shown as grey spheres. Images generated using Chimera (Pettersen et al., 2004).

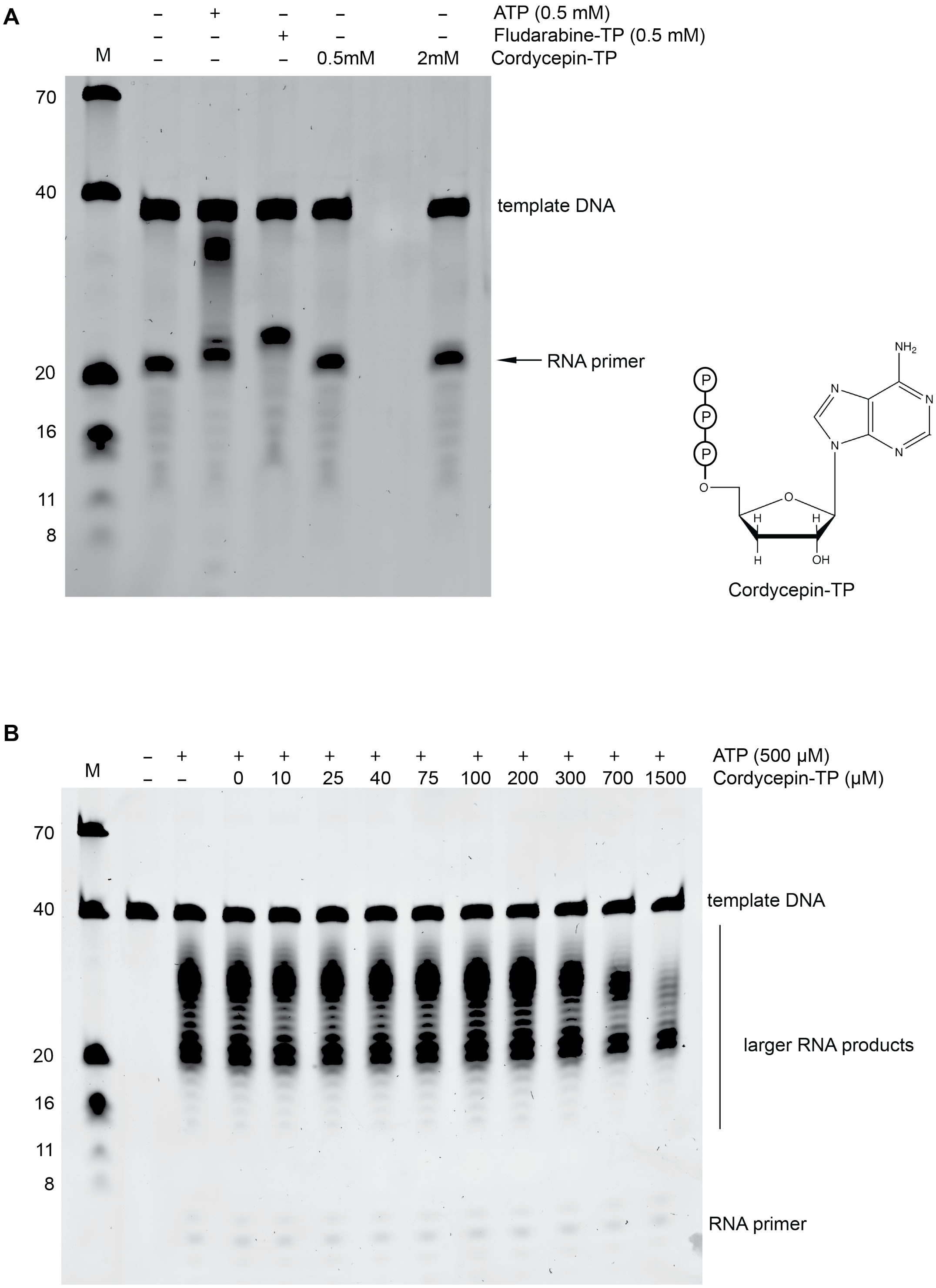
A
ATP (0.5 mM)
Fludarabine-TP (0.5 mM)
Cordycepin-TP
M
0.5mM
2mM
70
40
20
16
11
8
template DNA
RNA primer
Cordycepin-TP
B
ATP (500 µM)
Cordycepin-TP (µM)
0 10 25 40 75 100 200 300 700 1500
larger RNA products

**Supplementary figure 4**: Cordycepin-TP is poorly incorporated into an RNA primer and only inhibits primase acitivity at relatively high concentrations. (a) Denaturing gel showing incorporation of Cordycepin vs Fludarabine into an existing RNA primer. The template comprised a 38-mer DNA (5′-$T_{20}$CCAGAGAGCGCCCAAACG) annealed to an 18-mer RNA (5′-CGUUUGGGCGCUCUCUGG). Reactions contained 0.5 μM template, 0.5 μM primase, 10 mM $Mg(OAc)_2$ and 500 μM ATP, Fludarabine-TP or Cordycepin-TP. Reactions were incubated at 37 °C for 30 minutes, and the products were analysed by denaturing polyacrylamide gel electrophoresis. (b) Denaturing gel showing the effect of Cordycepin-TP on RNA primer synthesis. Reactions contained 0.5 μM polydT40 ssDNA template, 0.5 μM primase, 500 μM ATP, 10mM $Mg(OAc)_2$ and the indicated concentration of Cordycepin-TP. Products were incubated and analysed as in (a). All gels were post-stained with Sybr Gold. M = marker.

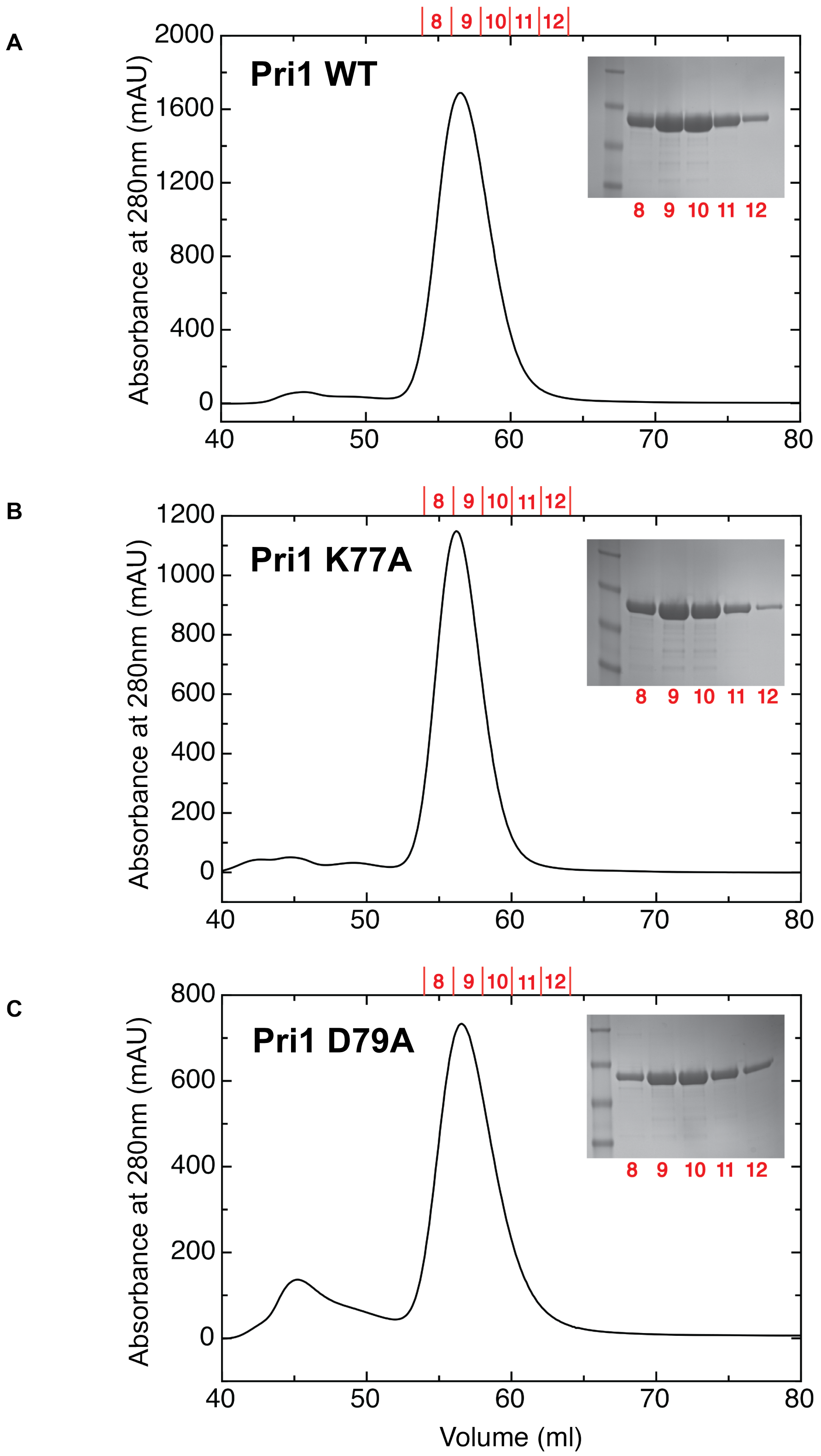
A
Pri1 WT
B
Pri1 K77A
C
Pri1 D79A
8 9 10 11 12
Absorbance at 280nm (mAU)
Volume (ml)
0
400
800
1200
1600
2000
0
200
400
600
800
1000
1200
40
50
60
70
80

**Supplementary figure 5**: Analysis of the D79A and K77A Pri1 point mutants by size exclusion chromatography. Elution profiles from a S75 16/60 gel filtration column for Pri1 (a) wild-type, (b) K77A and (c) D79A. Coomassie-stained SDS-PAGE gels of the eluted fractions are shown as inset panels.

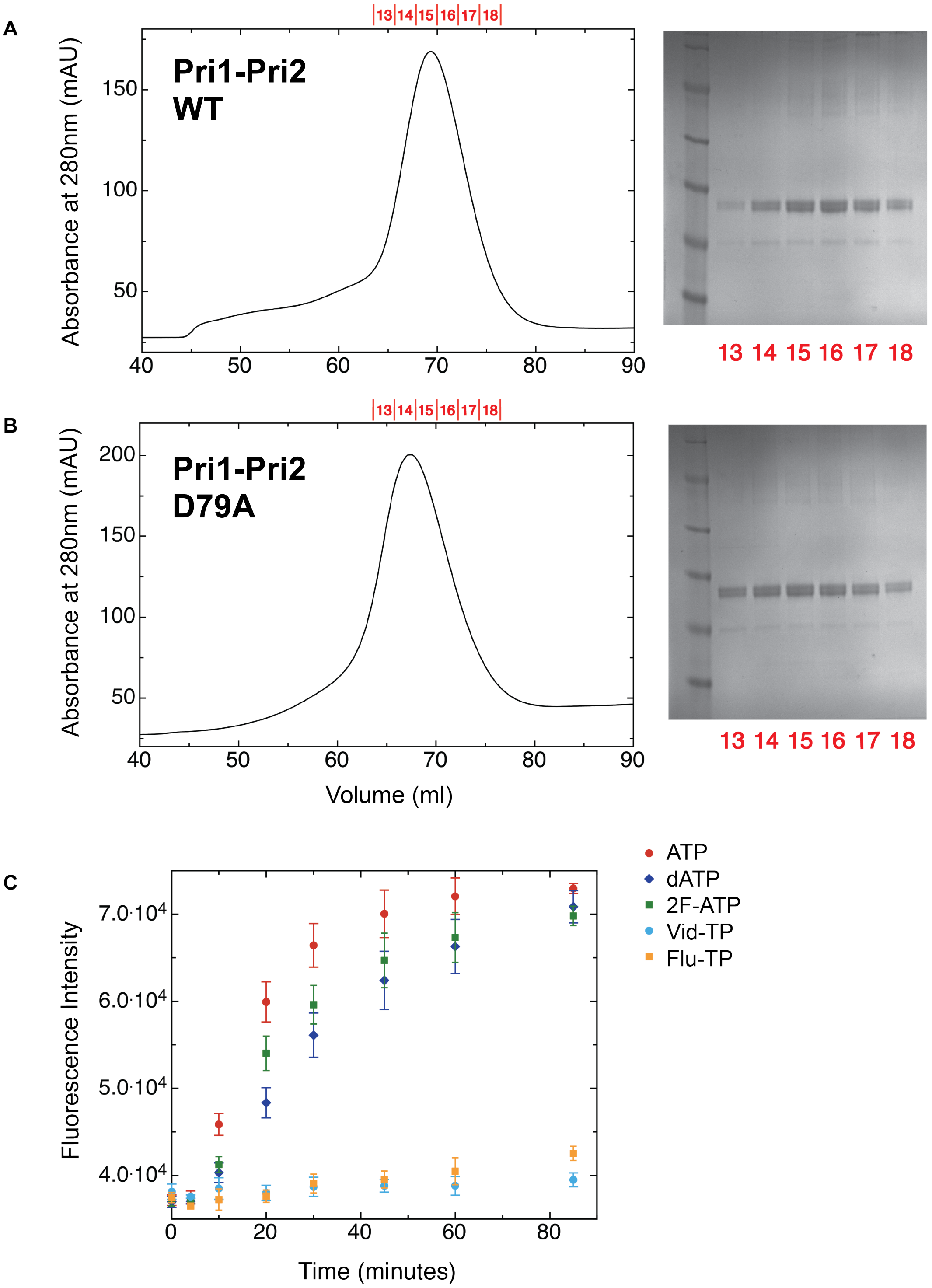
A
B
C
Pri1-Pri2
WT
Pri1-Pri2
D79A
Absorbance at 280nm (mAU)
Volume (ml)
13 14 15 16 17 18
Fluorescence Intensity
Time (minutes)
ATP
dATP
2F-ATP
Vid-TP
Flu-TP

**Supplementary figure 6**: Analysis of the D79A Pri1-Pri2 point mutant

Elution profiles from a S200 16/60 gel filtration column for Pri1-Pri2 (a) wild-type and (b) D79A point mutant. Coomassie-stained SDS-PAGE gels of the eluted fractions are shown as inset panels. Due to size similarities, Pri1(1-420) and Pri2(1-462) migrate as a close doublet. (c) Fluorescence-based RNA primer synthesis assay on a ssDNA template (5′-GTTGTCCATTATGTCCTACCTCGTGCTCCT) in the presence of $Mn^{2+}$ ions and equimolar concentrations of nucleotide (20 μM each NTP) and the indicated nucleotide analogue (20 μM). Each data point represents the mean ± s.d. (n=3). Curves are coloured as follows: ATP (red), dATP (navy), 2F-ATP (green), Vidarabine-TP (light blue), Fludarabine-TP (orange).

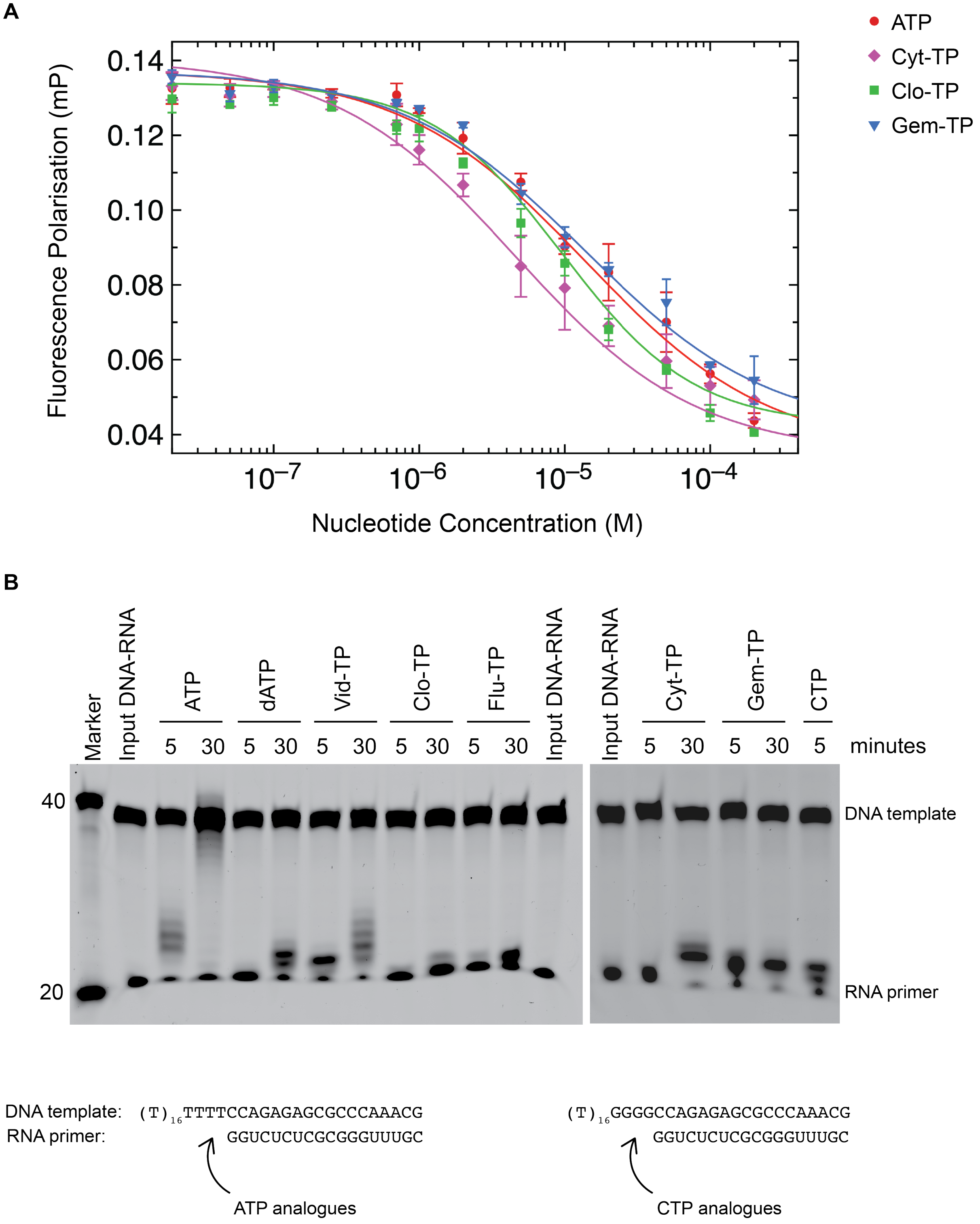
A
ATP
Cyt-TP
Clo-TP
Gem-TP
Fluorescence Polarisation (mP)
0.14
0.12
0.10
0.08
0.06
0.04
$10^{-7}$
$10^{-6}$
$10^{-5}$
$10^{-4}$
Nucleotide Concentration (M)
B
Marker
Input DNA-RNA
ATP
dATP
Vid-TP
Clo-TP
Flu-TP
Input DNA-RNA
Input DNA-RNA
Cyt-TP
Gem-TP
CTP
5 30 5 30 5 30 5 30 5 30
5 30 5 30 5
minutes
40
20
DNA template
RNA primer
DNA template: $(T)_{16}$TTTTCCAGAGAGCGCCCAAACG
RNA primer: GGUCUCUCGCGGGUUUGC
ATP analogues
$(T)_{16}$GGGGCCAGAGAGCGCCCAAACG
GGUCUCUCGCGGGUUUGC
CTP analogues

**Supplementary figure 7**: Analysis of the effect of Cytarabine-TP, Clofarabine-TP and Gemcitabine-TP on RNA primer synthesis. (a) FP-based nucleotide binding experiment in which Pri1 was pre-bound to 6FAM-labelled ATP then challenged with increasing concentrations of the indicated nucleotide (ATP) or nucleotide analogue (Cyt-TP: Cytarabine triphosphate, Gem-TP: Gemcitabine triphosphate, Clo-TP: Clofarabine triphosphate). Each data point represents the mean ± s.d. (n=3, except Cyt-TP for which n=6). (b) Denaturing gel showing the incorporation of nucleotide analogues into an existing RNA primer. The template comprised a 38-mer DNA template (5′-$T_{16}$TTTTCCAGAGAGCGCCCAAACG) annealed to an 18-mer RNA (5′-CGUUUGGGCGCUCUCUGG). For the cytosine analogues (Cytarabine-TP and Gemcitabine-TP) the DNA template was instead 5′-$T_{16}$GGGGCCAGAGAGCGCCCAAACG. Reactions contained 0.5 μM annealed DNA-RNA template, 0.5 μM primase, 10mM $Mg(OAc)_2$, and 500 μM ATP, dATP, Vid-TP, Clo-TP, Flu-TP, Cyt-TP or Gem-TP. Reactions were allowed to proceed for 5 or 30 minutes at 37 °C before quenching and subsequent separation of the products on an 18% polyacrylamide-urea gel. Gels were post-stained with Sybr Gold. M = marker.

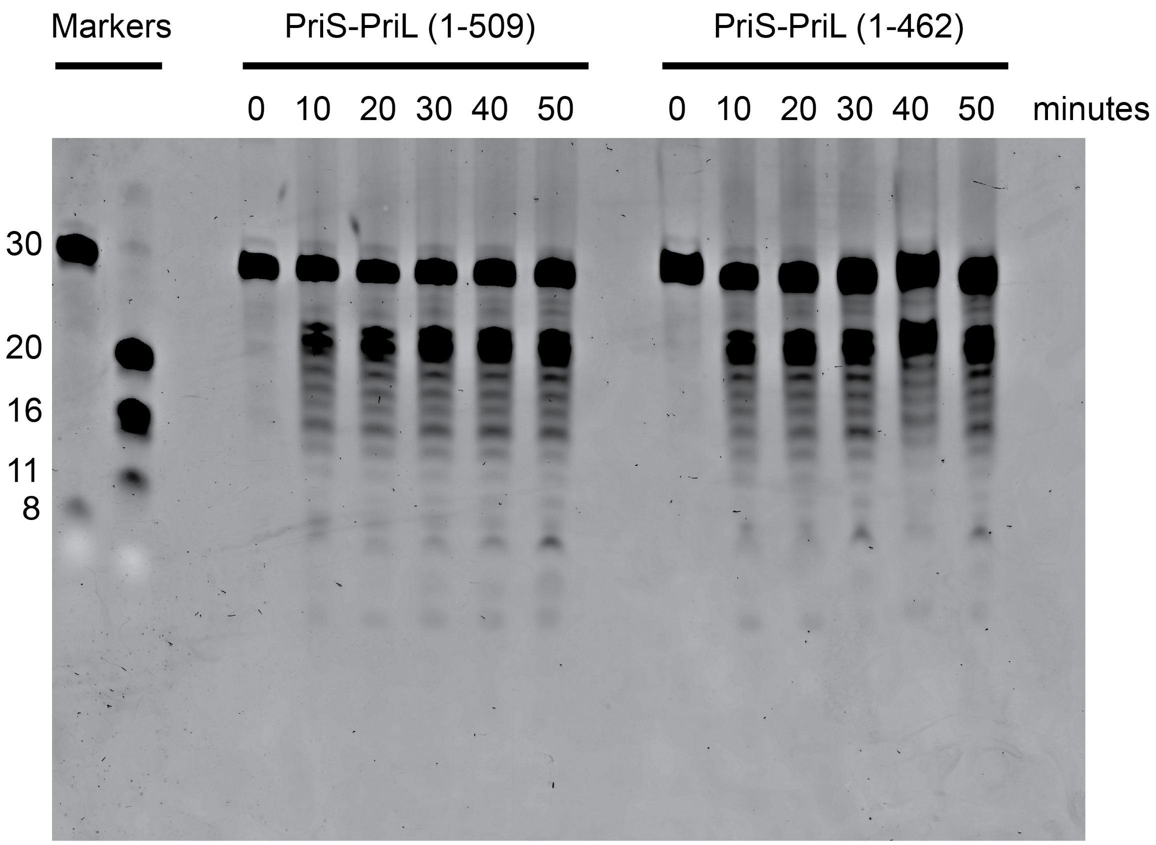


**Supplementary figure 8**: Pri1-Pri2(1-509) and Pri1-Pri2(1-462) display similar levels of activity Denaturing gel showing the time-course of RNA primer synthesis by Pri1(1-420)-Pri2(1-509) (left) and Pri1(1-420)-Pri2(1-462) (right). Primer products were visualized by post-staining the urea-polyacrylamide gel with Sybr Gold.
